# Toward optimal fingerprint indexing for large scale genomics

**DOI:** 10.1101/2021.11.04.467355

**Authors:** Clément Agret, Bastien Cazaux, Antoine Limasset

## Abstract

**Motivation:** To keep up with the scale of genomic databases, several methods rely on local sensitive hashing methods to efficiently find potential matches within large genome collections. Existing solutions rely on Minhash or Hyperloglog fingerprints and require reading the whole index to perform a query. Such solutions can not be considered scalable with the growing amount of documents to index.

**Results:** We present NIQKI, a novel structure with well-designed fingerprints that lead to theoretical and practical query time improvements, outperforming state-of-the-art by orders of magnitude. Our contribution is threefold. First, we generalize the concept of Hyperminhash fingerprints in (h,m)-HMH fingerprints that can be tuned to present the lowest false positive rate given the expected sub-sampling applied. Second, we provide a structure able to index any kind of fingerprints based on inverted indexes that provide optimal queries, namely linear with the size of the output. Third, we implemented these approaches in a tool dubbed NIQKI that can index and calculate pairwise distances for over one million bacterial genomes from GenBank in a few days on a small cluster. We show that our approach can be orders of magnitude faster than state-of-the-art with comparable precision. We believe this approach can lead to tremendous improvements, allowing fast queries and scaling on extensive genomic databases.

**Availability and implementation:** We wrote the NIQKI index as an open-source C++ library under the AGPL3 license available at https://github.com/Malfoy/NIQKI. It is designed as a user-friendly tool and comes along with usage samples.

**2012 ACM Subject Classification:** Applied computing → Bioinformatics

**Digital Object Identifier:** 10.4230/LIPIcs.WABI.2022.25

## Introduction

Historically, genomic databases such as GenBank are growing exponentially^3^. Lower sequencing costs and required investment, broader access to sequencing technologies, and breakthrough in genome assembly practices will surely fuel the explosion of available genomes in the near future. While those data are still widely unprocessed, the ability to explore and delve into such rich archives presents countless applications [4, 1]. Allowing to query such databases at a reasonable cost is a growing research subject [3, 13, 15, 16]. To avoid relying on computation-intensive steps, an efficient way to approximate the similarity of two sequences is to represent them as a set of *k*-mer (sub-word of length *k*) and compare their *k*-mer contents. Namely, the fraction of shared *k*-mer that corresponds to the Jaccard index is a good proxy for Average Nucleotide Identity [14] between genomes. Given the scale of such collections, indexing complete genomes quickly become prohibitively expensive. When searching for broad matches between large genomic sequences such as genomes, dimension reduction techniques can be used to reduce the computational burden.

Minhash [5] is a very resource-efficient technique able to estimate the Jaccard index between two sets. Minhash constructs a sketch of *S* fingerprints chosen from the hashed elements of a given set of size *N* (e.g., selecting the smallest hash values). The attractive property of such sketches is that the Jaccard index between them approximates the actual Jaccard index of the two sets they represent. This way, the Jaccard index can be estimated by comparing two sketches of size *S* that can be orders of magnitude smaller than the actual cardinality of the indexed sets.

Three principal variants of Minhash have been proposed.

### S hash functions

Each element is hashed using *S* distinct hash functions *h*_1_ … *h*_*S*_, the sketch is composed of the smallest hash output by each hash function. The sketches can be compared in Θ(*S*) and constructing sketches is Θ(*N*.*S*)

### S minimal values

Each element is hashed using a single hash function *h*. The sketch comprises the *S* smallest hashes output by *h*. The sketch construction is reduced to Θ(*N*. log(*S*)) but the sketch comparison is Θ(*S*. log(*S*))

### S partitions

Each element is hashed using a single hash function *h*. The hash values are split into *S* partitions according to their first bits. The sketch is composed of the minimal element of each partition. The sketch construction is reduced to Θ(*N*) and the sketch comparison to Θ(*S*).

While *S* partition seems optimal, some partitions can be empty if no hashes start with a given prefix. Some process dubbed densification [17] aims to cope with this problem by using other partition data to fill empty partitions.

The other parameter of a Minhash sketch is the size of each stored hash. Usually, a “large” hash size (e.g., 32 bits) is used to avoid the risk of collisions or saturation. Collisions occur when two different keys have the same hash value, leading to false-positive hits. Since the smallest hash values are kept, all fingerprints tend to zero. Once the zero value is reached, it can no longer change. We call such fingerprints saturated. A comparison of saturated sketches can report a large amount of false-positive hits.

However, since the minimal values are kept, the first bits of each fingerprint may contain a large amount of low-informative zeroes. The b-bit Minhash [9] variation takes advantage of this observation and only keeps the b lowest bits to reduce the sketches sizes by reducing the fingerprints sizes. It can either improve the memory footprint of sketches or their accuracy by increasing the number of fingerprints used for the same amount of memory.

Those techniques were successfully applied in bioinformatics to index and compare genomes databases. Mash [14] implemented *S* minimal values and Bindash [19] implement *S* partitions, b-bit Minhash with densification. Such works showed that genomes and whole sequencing datasets could be precisely compared by using sketches orders of magnitude smaller than their amount of distinct *k*-mers leading to tremendous performance improvement when working at a large scale.

An alternative to Minhash fingerprints is Hyperloglog [6] fingerprints. Instead of storing the whole hash, Hyperloglog fingerprints store its log value (e.g., the position of the leading 1). The interest in such fingerprints is twofold. They are tiny: a 64bits hash needs a 6bits Hyperloglog fingerprint (since 2^6^ = 64) and is as hard to saturate as their associated hashes. Six bits seem enough to cover most real-world cases in practice, as 64bit hashes seem unlikely to be saturated. The downside is that their collision rate is very high. This fact can be leveraged by using a large number of fingerprints. Dashing [2] showed that large *S* partitions sketches of Hyperloglog fingerprints could lead to very precise estimations of Jaccard index between genomes.

Interestingly, another approach tried to benefit from the trailing zeroes by combining the b-bit Minhash and Hyperloglog ideas. Hyperminhash [18] builds a fingerprint from a given hash using 6 bits to encode the first run of zero (which is equivalent to a Hyperloglog fingerprint) and combines them with the lowest 10 bits (which is equivalent to a b-bit Minhash fingerprint) to obtain 16 bits fingerprints. The interest of this sketch is to be very hard to saturate due to the Hyperloglog fingerprint and to limit the number of collisions with the additional Minhash bits. Such fingerprint can be seen as an efficient lossy compression of the hash value that represents more than 16 bits from the hash (if the run length of zeroes is longer than 6).

In bioinformatics, such fingerprint indexes have numerous applications such as very fast clustering or phylogeny construction from large genome collections, finding potentially related genomes from assembled contigs or sequences of interest, quantification of a given gene, strain, or species in databases etc... Their main feature is to find possible matches for a query sequence inside a database, acting as filters to focus only on relevant entries. They do so by efficiently estimating similarities between query and index sequences.

The main bottleneck of existing methods is that a query requires the queried sketch to be compared to all sketches in the database. The query is 𝒪(*N*.*S*) with *N* the number of indexed entries. This cannot be considered scalable regarding the exponential growth of available genomic resources. To cope with this algorithmic problem, we design a structure to perform queries in Θ(#*hits*) where #*hits* is the number of fingerprint hits between the queried sketch and the indexed sketches. Since the expected output of a query is a list of genomes identifier associated with a number of hits (or an approximation of the Jaccard index), we can consider our query time as close to optimal.

However, such a structure presents an overhead exponential with the fingerprint size. For this reason, we are highly attentive to using powerful but small fingerprint sizes. We generalize the Hyperminhash fingerprint and introduce (*h, m*)-Hyperminhash fingerprints that cover both Minhash and Hyperloglog cases and provide an analysis to select the best fingerprint for a given use case.

## Methods

### Index structure

To generalize the concept of indexing a fixed amount of fingerprints (either Hyperminhash, b-bit Minhash, or Hyperloglog), we call such a tuple of fingerprints a sketch. In the following, we consider bit-vectors of size *n* and integers between 0 and 2^*n*^ − 1 as equivalent. For the sake of clarity, log() will be used for log_2_() in this document.

Let *W, S*, and *N* be three integers, where *N* will be the number of sketches, *S* the size of the sketches, which is the number of fingerprints in each sketch, and *w* the size in bits of each fingerprint. More specifically, a *sketch s*_*i*_ of size *S* is a tuple of *S* fingerprints where a fingerprint is an element between 0 and 2^*W*^ − 1, i.e. *s*_*i*_ = *f*_*i*,1_ … *f*_*i,S*_ where ∀ *j* ∈ {1,…, *S*} *f*_*i,j*_ ∈ {0 − 2 ^*W*^ −1}. We present a new data structure to index a set *B* = {*s*_1_, …, *s*_*N*_} of *N* sketches where we want to optimize the time complexity of the query `find_hits(B, s′, m)` which corresponds to find all the sketches *s*_*i*_ = *f*_*i*,1_ … *f*_*i,S*_ of *B* such that 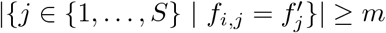 with 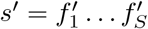 a sketch and *m* an integer.

Instead of indexing each sketch, we split the set of sketches *B* by column *j* between 1 and *S*. We denote by *P*_*j*_ the sequence of fingerprints *f*_1,*j*_ … *f*_*N,j*_. Unlike previous approaches, our scheme handles each column *P*_*j*_ independently by grouping the indices with the same fingerprint on this column. Indeed, for each column *j* between 1 and *S*, we index in a vector the sets 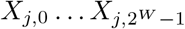 where for all *l* ∈ {0, …, 2^*W*^ − 1}, *X*_*j,l*_ = {*I* ∈ {1, …, *n*} | *f*_*i,j*_ = *l*}. 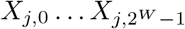 is a partition of {1, …, *N*}, our vector have 𝒪(*N* + 2^*W*^) elements (see Figure 1).

**Figure 1.**
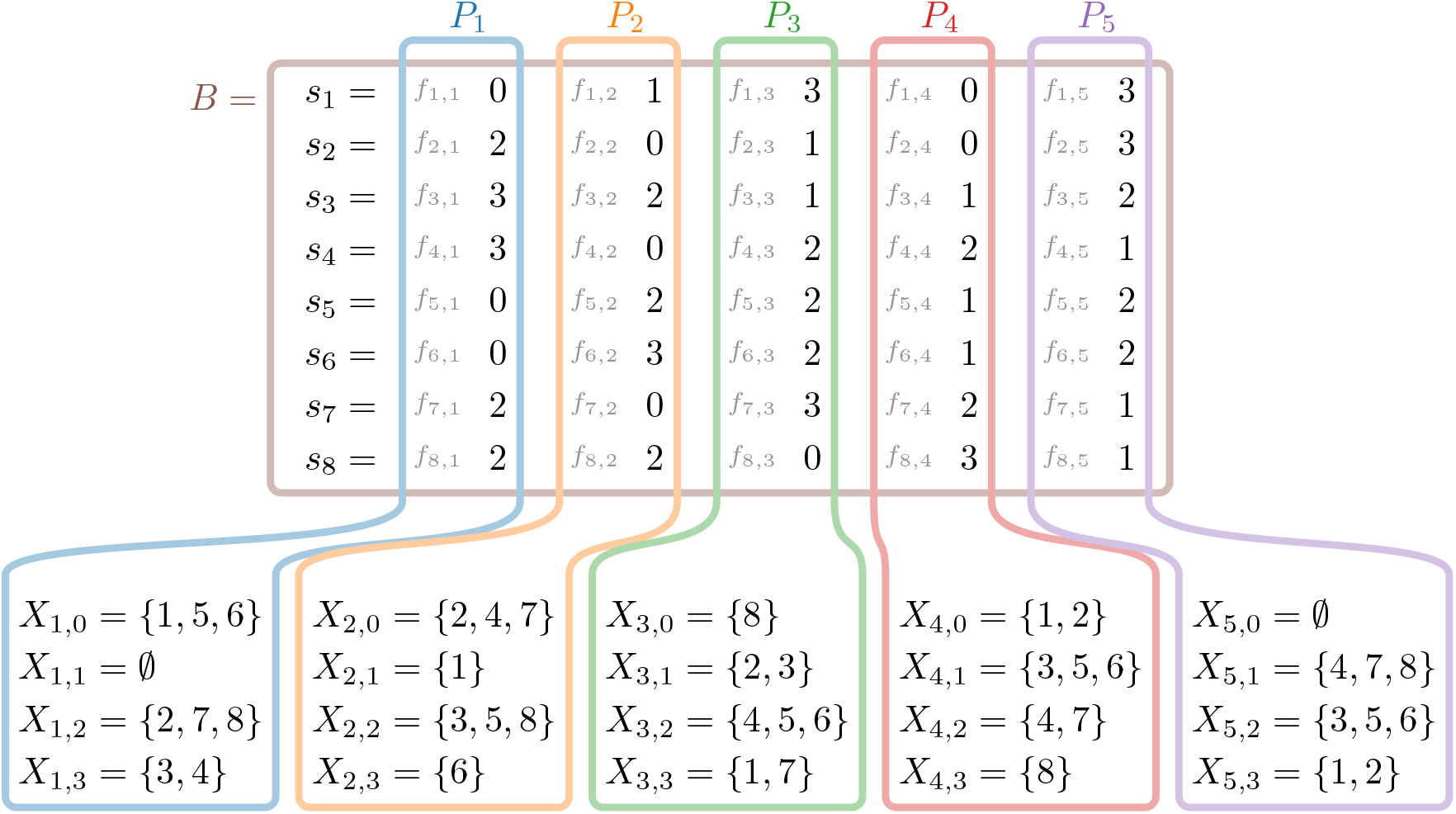
Example of processing partitions of 8 (*N*) sketches where each sketch has 5 (*S*) fingerprints of 2 (*W*) bits. *B* = {01303, 20103, 32112, 30221, 02212, 03212, 20321, 22031} is the set of sketches used as instance where the sketch *s*_1_ = 01303 corresponds to the tuple of the 5 fingerprints where the first fingerprint is 0, the second fingerprint is 1, the third fingerprint is 3 and so on. To represent the decomposition by column, we add specific color. For the first column (in blue) *P*_1_ corresponds to all the fingerprints seen as the first position of a sketch of *B* (the first partition) and *X*_1,*vf*_ contain the set of sketch identifiers (positions of the sketches in *B*) whose first fingerprint value is *vf* . We obtain *S* × 2^*W*^ = 20 sets *X*_*j,l*_ (1 ≤ *j* ≤ *S* and 0 ≤ *l* ≤ 2^*W*^ − 1) corresponding to the stored sets.

We argue that performing queries by partition presents several advantages. One advantage of this processing partition by column is the possibility to stop queries that can not find high-scoring matches. If we are looking for a minimum of *m* shared fingerprint among *S* to report a match, if after *P* _|*S*| −*j*_ no sketches share at least *m* − *j* fingerprints, that query can be stopped as no matches can reach the minimal score. If not using the densification technique, empty partitions could also be skipped.

As, for our case, a sketch represents a genome, and the fingerprints come from the set of *k*-mers of this genome, one possible mechanism presented in [10] inspired by [7] is to insert the *k*-mers of a given partition in a Bloom filter (or any set structure) to “protect” each partition from alien *k*-mers. If the selected *k*-mer is not in the partition set, the partition is skipped as we know that the *k*-mer was not inserted in the partition and that any fingerprint hit would be a false positive.

We use dense addressing to index our partitions in a trade-off favoring code simplicity and throughput over memory footprint. This way, our index has an overhead of Θ(*S*.2^*W*^) as we allocate *S*.2^*W*^ vectors to represent all possible values of each partition. Once the index created, inserting the *n*th sketch *s*_*n*_ = *f*_1_ … *f*_*S*_ is inserting *n* in the vector *f*_*i*_ + *i*.2^*w*^ . The inserted elements cost Θ(*N*. log(*N*)) space and each partition can be queried for `find_hits`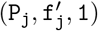in Θ(#*hits*) which corresponds to 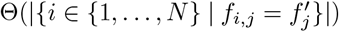. As `find_hits`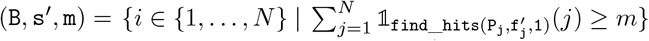, by computing all the `find_hits`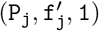, we can compute `find_hits (B, s′, m)`.

If this approach presents a costly overhead, its interest is that query a sketch performs at worst *S* “costly” random access. We argue that using relatively small, well-designed fingerprints, our scheme can perform swift queries with a reasonable memory footprint.

### Optimal Hyperminhash fingerprint

Since our proposed scheme presents an overhead exponential with *W*, we need to rely on very space-efficient fingerprints and make the most of each bit. When building a fingerprint database, the users tune the sketches’ size according to the expected genome sizes they index and the needed precision according to their application. This parameter also changes the fingerprint values as we expect each fingerprint to be the minimum across 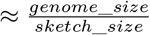 that we call the sub-sampling value. In the section, we argue that different sub-sampling values call for different fingerprints. This part presents a parametric fingerprint that generalizes Hyperminhash, b-bit minhash, and Hyperloglog. We give a method to fix this parameter to optimize the false positive rate for a given fingerprint size.

For a bitvector/hash *B* of size *n* and two integers *h* and *m* such that *h* + *m* = *W*, the (*h, m*)-hyperminhash fingerprint of *B*, denoted by `HMH`_*h,m*_(*B*), is the bitvector of size *h* + *m* where the *h* first bits correspond to the position to the first one on the prefix of *B* of size 2^*h*^ − 1 and the *m* last bits are the last *m* bits of *B*, i.e.

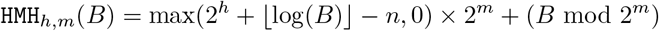

For a set *X* of bitvectors of size *n*, the (*h, m*)-hyperminhash fingerprint of *X* is the minimum value of the (*h, m*)-hyperminhash fingerprint of *B* for all the bitvectors *B* of *X*, i.e. `HMH`_*h,m*_(*X*) = min_*B*∈*X*_ `HMH`_*h,m*_(*B*).

Our goal is to find the pair *h, m* where *h* + *m* = *W* which gives us the most space-efficient fingerprint, i.e. the fingerprint which covers the most larger interval of values. One way to compute this space-efficiency is for a threshold *ε* of [0, 1], to compute *c*_*h,m*_ = *b*_*h,m*_ − *a*_*h,m*_ + 1 which is the size of the interval {*a*_*h,m*_, …, *b*_*h,m*_} where

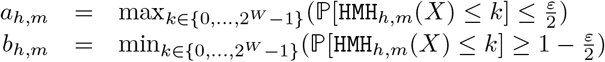

and thus ℙ [*a*_*h,m*_ `HMH`_*h,m*_(*X*) *b*_*h,m*_] ≤ 1 − *ε*.

To begin, if *n <* 2^*h*^ − 1 + *m*, some bits can be in both prefix and suffix of the fingerprint due to the overlap, and thus it is not space-efficient. Indeed, we can prove that the number of fingerprint values that are not reached is equal to 2^*m*^ × (2^*h*^ − 1 + *m* − *n*). For this reason, we take in all the following *n* ≥ 2^*h*^ − 1 + *m*.

For an integer *k* of {0, …, 2^*h*+*m*^}, we denote by *H*(*h, m, k*) the number of *B* ∈ {0, …, 2^*n*^ − 1} such that `HMH`_*h,m*_(*B*) = *k*. As (*h, m*)-hyperminhash is an increasing function because if *B* ≤ *B*′, we have `HMH`_*h,m*_(*B*) ≤ `HMH`_*h,m*_(*B*′), by counting, we can show that 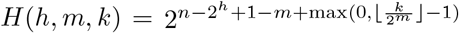. Indeed, each *B* of {0, …, 2^*n*^ − 1} such that `HMH`_*h,m*_ (*B*) = *k* has the suffix of length *m* and the prefix of length 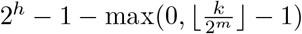 which are fixed.

As 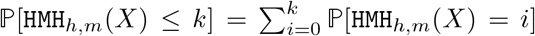, it is enougth to know the value of ℙ[`HMH`_*h,m*_(*X*) = *k*]. As 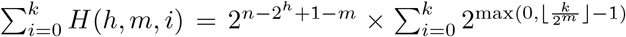, if *k* < 2^*m*^, 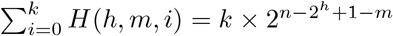. Otherwise,

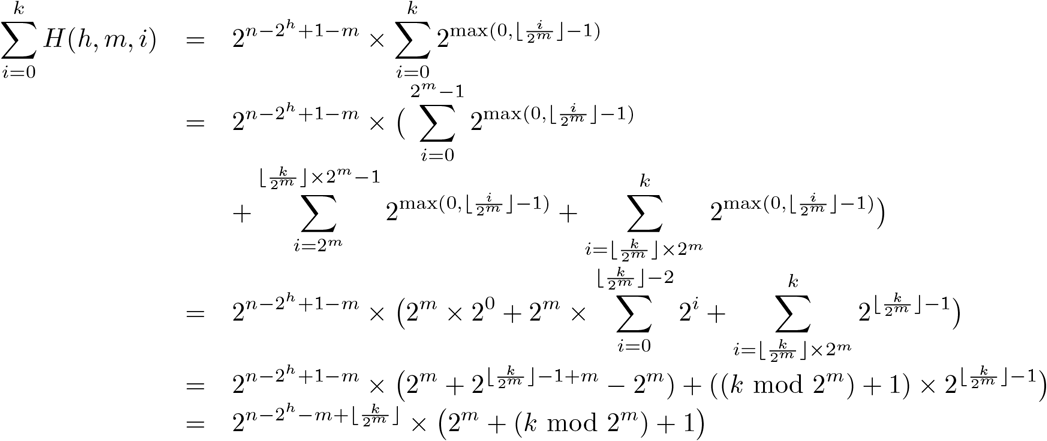

For the sake of completeness, we added all the calculations of 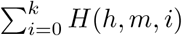, but the only thing one needs to remember is that it can be calculated in constant time. Besides, we have

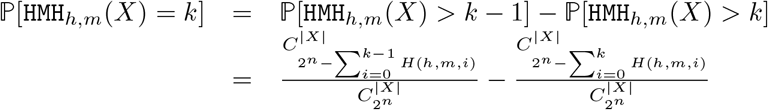

We can extend this formal definition to the probability of interval {*a*, …, *b*}, i.e.

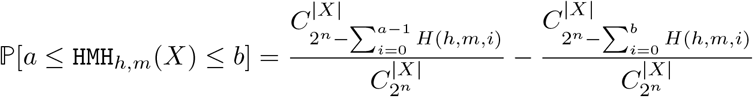

By approximating the value of ℙ[`HMH`_*h,m*_(*X*) ≤ *k*] by 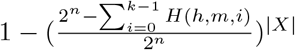, we can give an approximation of *a*_*h,m*_ and *b*_*h,m*_ for a threshold *ε* of [0, 1] in 𝒪 (1):

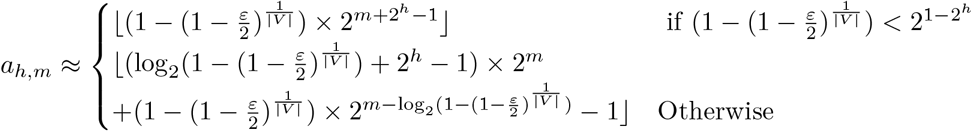

and

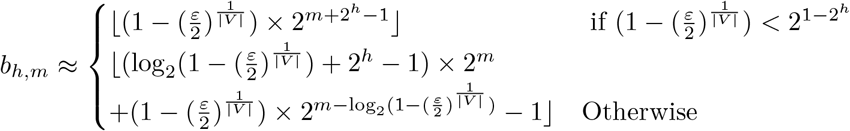

As shown in Figure 2a, the choice of the good fingerprint depends on the size of the sub-sampling. By dichotomic search, we can compute each exact value *c*_*h,m*_ in 𝒪 (|*X*| × *W*) and thus we can find the pair *h, m* which maximizes *c*_*h,m*_ in 𝒪 (|*X*| × *W* ^2^). In practice, we use the approximation value of *c*_*h,m*_ to compute an approximate optimal pair *h, m* in 𝒪(*W*).

**Figure 2.**
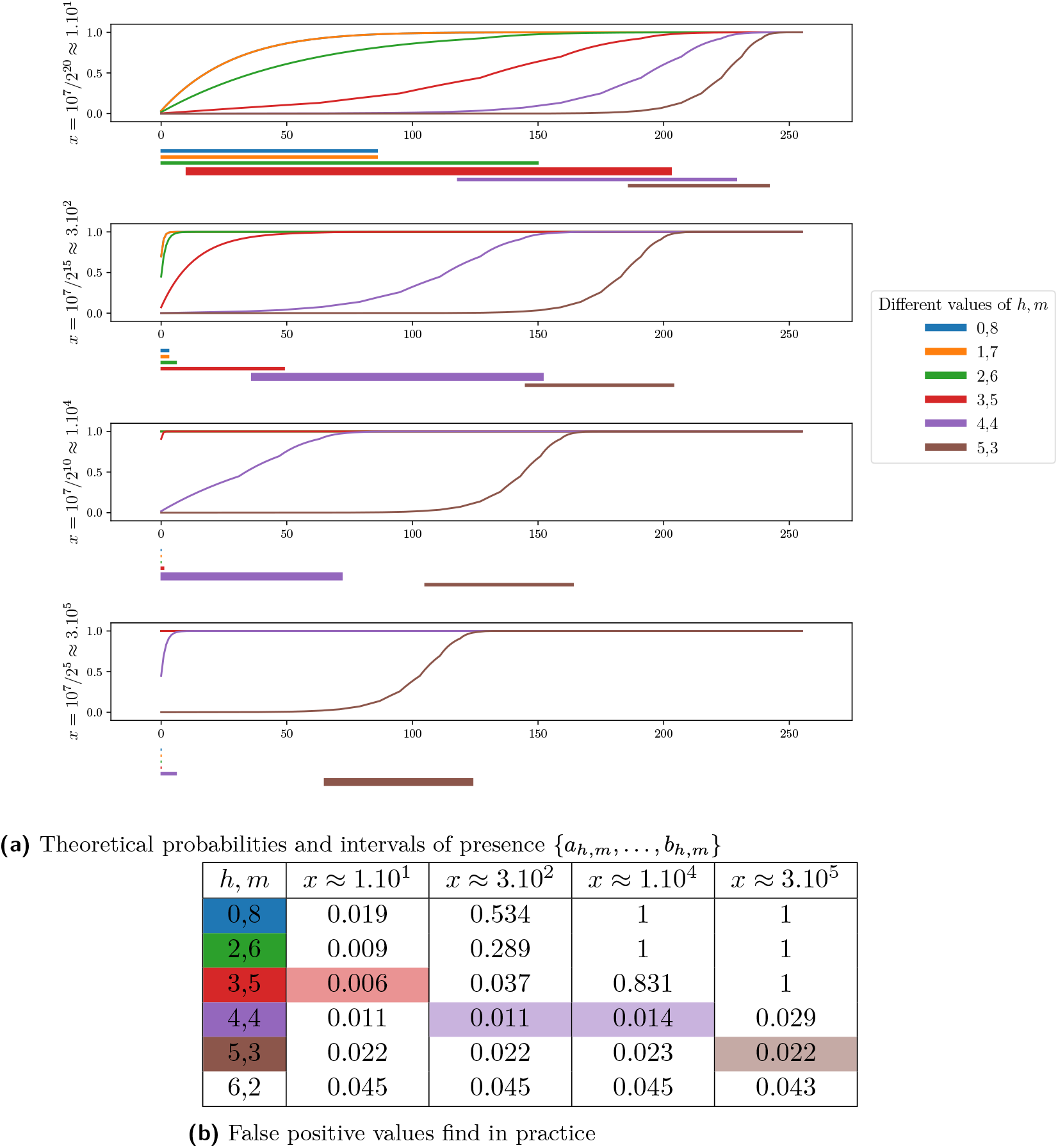
Correspondence between fingerprint with the theoretical maximal interval {*a*_*h,m*_, …, *b*_*h,m*_} and the fingerprint with the minimum number of false positive value in practice where *n* = 256, *W* = 8 and the number of *k*-mers is 10^7^ for different values of *S* ∈{2^5^, 2^10^, 2^15^, 2^20^} . 2a shows the different values of ℙ [`HMH`_*h,m*_(*X*) ≤ *k*] for all the *k* ∈ {0, …, 2^*W*^ − 1 = 255} and for all pairs *h, m* such that *n* ≥ 2^*h*^ − 1 + *m* and all values of *x* = number of *k*-mers*/S*. Under each plot, we add a rectangle for each interval {*a*_*h,m*_, …, *b*_*h,m*_} with the corresponding color. 2b gives the different values of false positive depending of a pair *h, m* and a subsampling size *x*.

The correspondence between the theoretical optimal fingerprint and the fingerprint with the minimum false positive values in practice justifies the relevance of our study (see Figure 2).

### Implementation details

Our proof of concept is dubbed NIQKI (stand for Next Index to Query *k*-mer Intersection), is open-source and available on Github https://github.com/Malfoy/NIQKI. In this section, we detail some practical aspects of this implementation.

Our NIQKI index is technically able to index any kind of fingerprint. In practice, from a given fingerprint size (12 by default), we use the best possible (h,m)HMH fingerprint when the user specifies an expected genome size or a regular Hyperminhash fingerprint if the expert user precise the size of the Hyperloglog fingerprint. We use a default Hyperloglog fingerprint of four if no input is given. We implemented the state-of-the-art densification technique [11] to take care of empty bins.

NIQKI is written in C++ and parallelized with straightforward OpenMP instructions. Each thread inserts or queries a set of sequences independently. A mutex array protects our vectors from double writing for insertion. We chose to use 64-bit hashes for performance purposes as they provide a low collision chance at the hash level in practice while using efficient 64 bits integers. For the same reason, we use a Xorshift [12] hash function are they are found good enough in practice while being incredibly cheap to compute. The sketch sizes are necessarily a power of two for code simplicity and performance aspects. Like Dashing, we ask the user to enter log_2_(*S*) instead of the actual sketch size.

Our implementation can take any classical sequence format as input: fastq, fasta, or multiline fasta files gzipped or not. The input is either a file of files where each file is an entry to index (or query) or a file where each sequence is a separate entry. We also provide a way to download genomes directly from NCBI from an accession list (that can be generated following NCBI instruction^4^). The corresponding sequences are downloaded and directly inserted into the index in a streaming fashion without any disk operation.

Indexes can be dumped and loaded from the disk for later use. This allows users to keep a small index on a disk instead of a large file collection and avoid reading all files to reload a given index.

Our implementation delivers a gzip-compressed sparse output to limit disk usage by default. We provide a pretty printer to parse such a file and an option to directly output (larger) human-readable output.

A common use of such a genome index is to compute all pairwise distances between them by comparing the index with itself. An efficient way is to check every vector of our index and increment the score of each genome pair present in the vector. While very fast in theory, this technique requires storing the whole score matrix of size *N* ^2^ in memory to be efficient. Henceforth this behavior can be interesting when a large amount of memory is available or when the number of genomes is not too large. Optimization relying on buffer keeping the matrix on disk can dissipate this memory footprint problem at the expense of several passes on the index. In the following benchmarks, the pairwise distances are computed without using the fact that the input is the indexed data itself. Competitors use this information to avoid reading the input file several times or computing the lower-left side of the symmetrical matrix.

## Results

### Fingerprint impact

As we previously evaluated the impact of the size of the Hyperloglog fingerprint, we now aim to evaluate the impact of the fingerprint size itself. The fingerprint size can be seen as the main parameter of our index and the amount of fingerprint used because our index has a *θ*(*S*.2^*W*^) memory overhead. More importantly, the fingerprint size will impact the amount of false-positive hits where two distinct hashes in compared sketches get the same fingerprint. Such false-positive hits over-estimate the similarity and may negatively impact downstream results by reporting irrelevant matches. To access the false positives found in practice, we created a synthetic dataset of one thousand randomly generated genomes of 10 million bases that share no *k*-mers. We computed the pairwise distances between those “genomes” and counted the number of hits between unrelated genomes as false positives. We computed sketches of size 4,096 from each ten megabases synthetic genome with fingerprint sizes varying from 4 to 18 with a constant Hyperloglog fingerprint size equal to four. We present the false-positive rates found in practice and the memory used by our index for such an experiment using various fingerprint sizes in Figure 3. As expected, we observe an exponential decrease in the false positive rate as we increase the fingerprint size. We observe that a small fingerprint size can obtain a shallow false-positive rate. From *W* = 12 the rate is below 10^−3^ and from *W* = 15 below 10^−4^.

**Figure 3.**
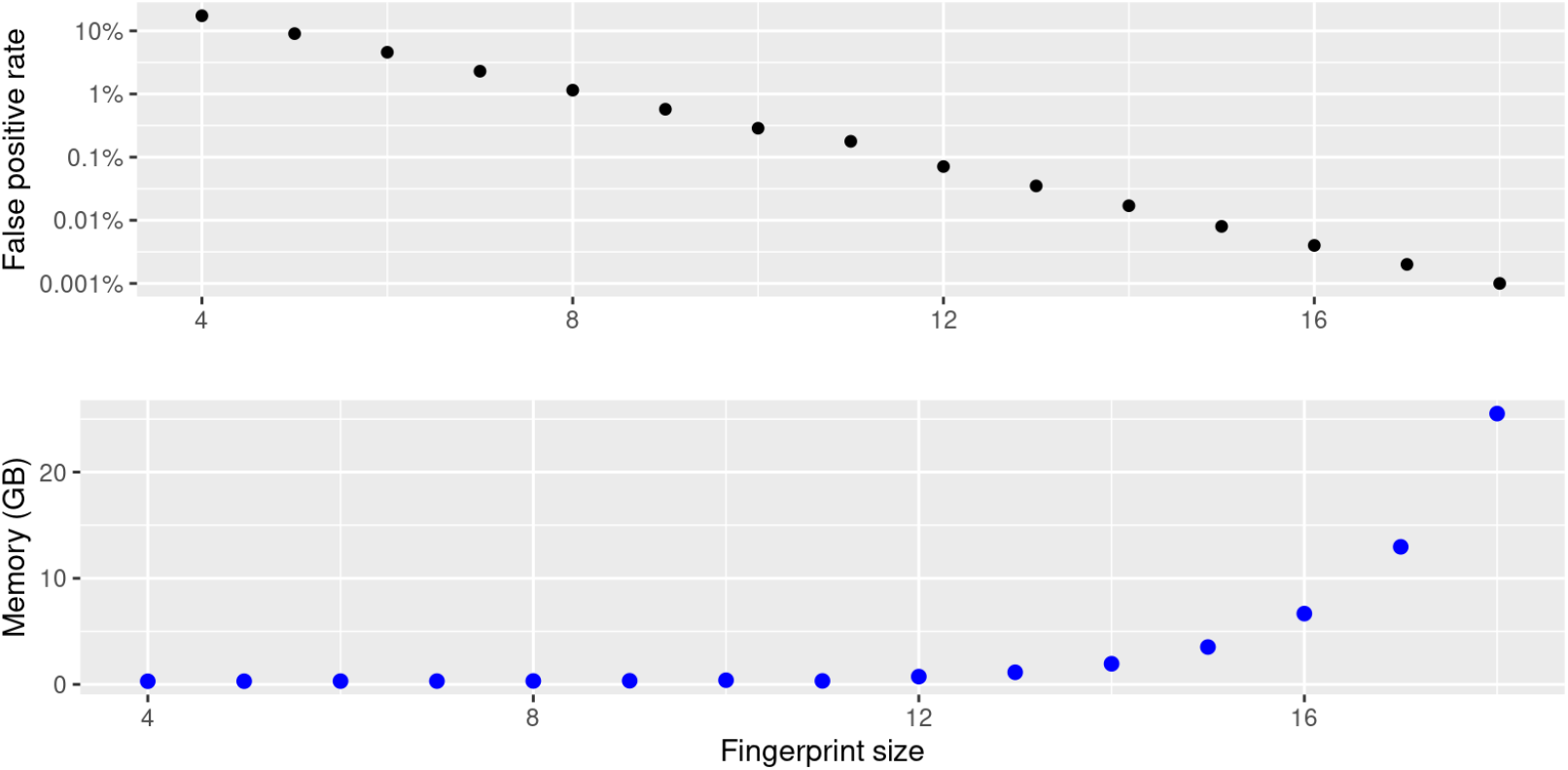
Impact of fingerprint size on false-positive rate and memory usage when indexing and comparing one thousand synthetic genomes of 10 megabases against themselves. Here the Hyperloglog fingerprint is kept constant (*H* = 4) and 4,096 fingerprint are used per genome.

### Accuracy analysis

To further analyze our approach’s precision, we compared the result of NIQKI with state-of-the-art on a real dataset composed of one thousand bacterial genomes from Refseq. In this experiment, we computed the pairwise distances between all genomes with our approach and compared those results with Dashing using many fingerprints (1 million) as the ground truth. In a first experiment reported in Figure 4 we plot the correlation between NIQKI and Dashing estimations using different amounts of fingerprint (namely *S* = 65536,*S* = 4096 and *S* = 256) while using a constant fingerprint size (*W* = 12). As described in the theoretical analysis of the related studies, a smaller amount of fingerprints results in larger error bounds. We observe that NIQKI obtain a strong correlation with Dashing output even with a very low amount of fingerprint. Even small sketches can lead to rough estimations that can be improved using larger sketches where the error bound can be tiny, as in the standard approaches.

**Figure 4.**
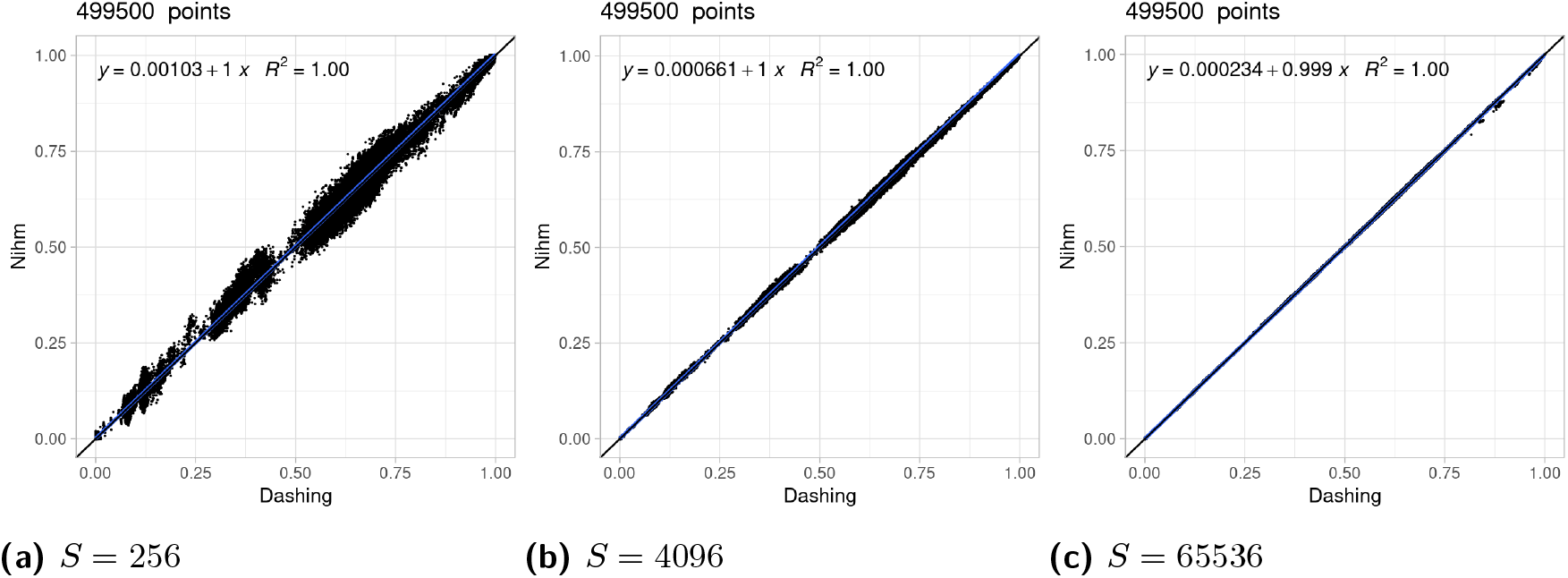
Impact of increasing sketch sizes. We display the correlation between NIQKI with varying sketch sizes and Dashing using one million fingerprints on one thousand bacterial genomes from RefSeq. Here the fingerprint size is kept constant (*W* = 12).

In a second experiment reported in figure5 we used different fingerprint sizes (namely *W* = 8,*W* = 6 and *W* = 4) with a constant fingerprint number(*S* = 65536).

Due to the large number of fingerprints, we observe very small error bounds, but due to the false positive rate, we observe that all indices are overestimated. The expected overestimation is ≈ (1 − *J*) ∗ *FP*_*Rate*_ where *J* is the Jaccard index between the two sequences, and the observed pattern goes accordingly with this projection.

This experiment shows that using fingerprints as small as eight bits seems reasonable in practice for such an index. The false positive rate could still impact the results when dealing with a low Jaccard index, and a larger fingerprint should be used. We fixed the default fingerprint size to 12 as it provides very low false positives and a reasonable memory overhead.

### Performance analysis

This section displays how our index can scale on synthetic and real datasets compared to the two most used tools of the state-of-the-art Mash and Dashing. All experiments were performed on a single cluster node running with Intel(R) Xeon(R) CPU E5-2420 @1.90GHz with 192GB of RAM and Ubuntu 16.04. with a timeout of 48hours. Bindash was not included in our benchmark because it cannot compute a distance matrix from a file of files directly.

All state-of-the-art tools present a 𝒪(*S*.*N*) query time (or 𝒪(*S*. log(*S*)).*N*) if S minimum Minhash is used) as a query sketch have to be compared to each indexed sketch. In contrast, our index presents a query time linear with the number of hits. It can be 𝒪(*S*.*N*) in the worst case where all genomes are identical and can be (*S*) in the best-case scenario where an alien entry is entirely dissimilar from the indexed genomes (ignoring false positives).

To show those two regimes, we perform a first “idealistic” benchmark with randomly generated genomes sharing no *k*-mers that display the best case of our index. We compare those results to a real database composed of GenBank genomes in a second experiment. We downloaded all Genbank bacterial assemblies (1,042,611) and randomly selected genomes from this pool to build such an index. We want to point out that such a database is highly redundant. For instance, it contains 142,568 assemblies of the Escherichia coli organism. It constitutes a stress case for our index as each query associated with Escherichia coli should report a large part of the indexed genomes among its matches.

In Figure 6 we ran Dashing, Mash, and NIQKI to compute pairwise distance on randomly generated genome collection of growing sizes and report the mean CPU time per entry and RAM usage. Because of the quadratic aspect of computing all pairwise distances, we report the mean time required per entry by dividing the total running time by the amount of entry to improve results readability. To provide a fair comparison, we choose parameters such as each tool to allocate the same memory per entry. Mash used 65,536 32 bits Minhash fingerprints, Dashing used 262,144 8 bits Hyperloglog signatures, and NIQKI used 65,536 32 bits genome identifiers and the default fingerprint (*W* = 12). We observe that both Mash and Dashing runtime grow following an expected quadratic evolution as computing all pairwise distances needs Θ(*S*.*N* ^2^). The growth runtime of NIQKI is roughly linear and can become orders of magnitude faster than state-of-the-art.

**Figure 5.**
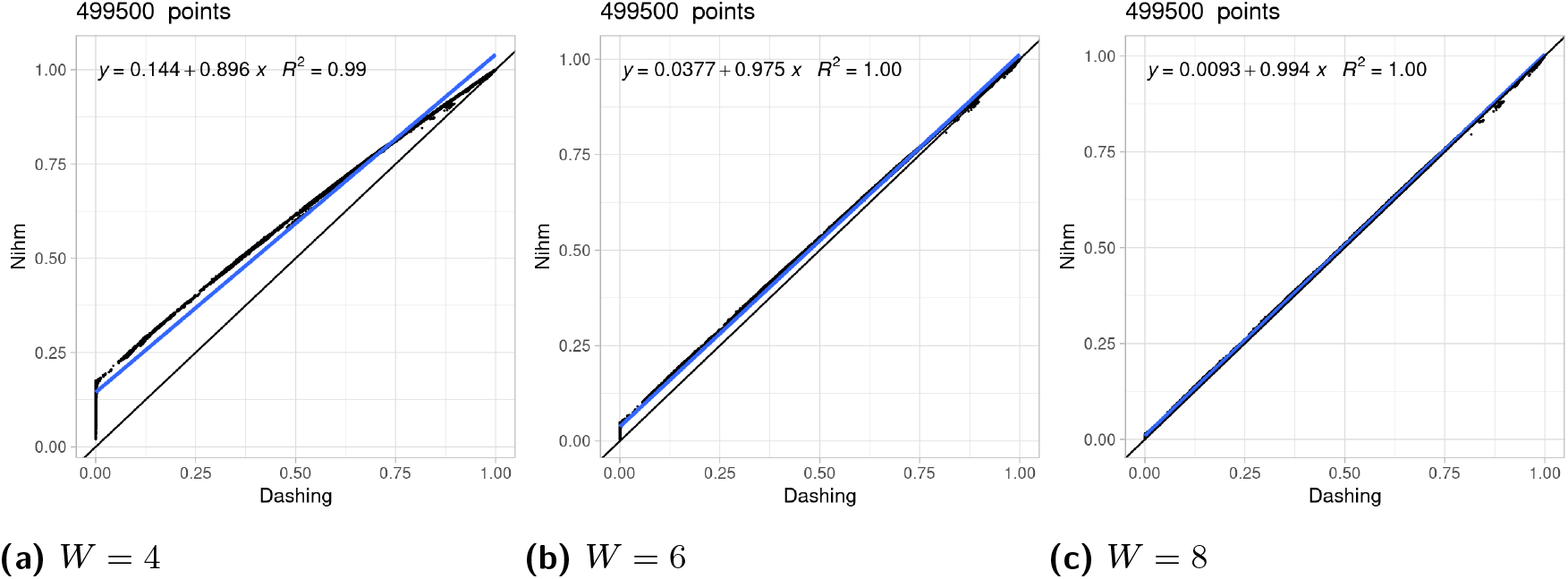
Impact of increasing fingerprint sizes. Correlation between NIQKI with varying fingerprint size and Dashing using one million fingerprints on one thousand bacterial genomes from RefSeq. Here the sketch size is kept constant (*S* = 65536).

**Figure 6.**
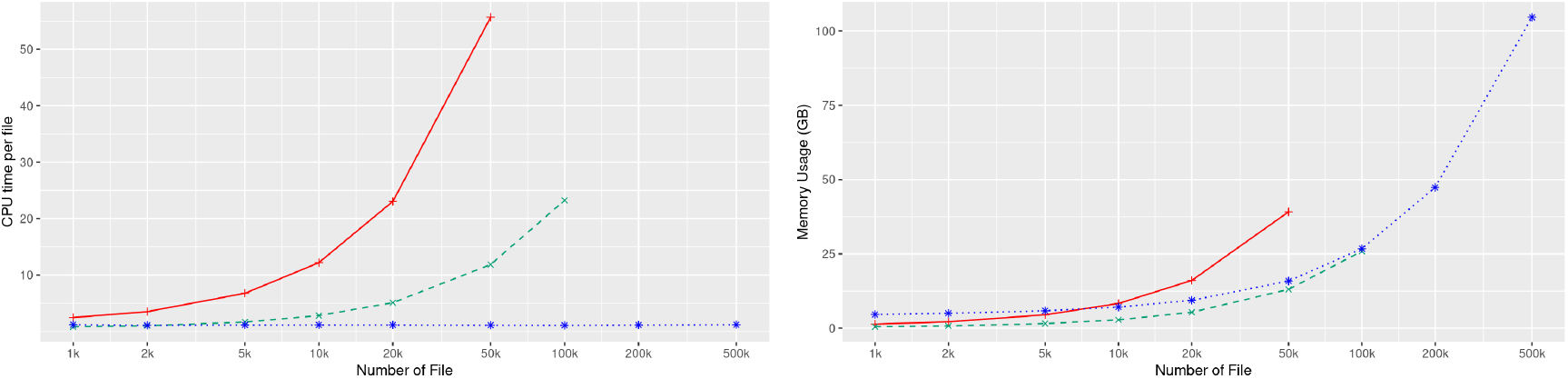
Benchmark on synthetic genomes. We report the CPU time divided by the number of entry and the total memory footprint for various collection sizes. Mash is plot in red, Dashing in green and NIQKI in blue.

In Figure 7 we perform the same experiment on a realistic database constituted from bacterial GenBank genomes. We observe that Mash and Dashing deliver similar performances on synthetic and real datasets. The differences are due to the fact that the mean size of real bacterial genomes is around five megabases, while the generated genomes are ten megabases long. NIQKI results on small databases are similar to the synthetic cases, but a superlinear growth can be observed on the largest collections. However, despite this observation NIQKI can still be orders of magnitude faster than state-of-the-art tools while using a comparable amount of memory. For example, the largest Mash experiment used 773 CPU hours on 50k genomes where NIQKI used only 15 CPU hours. The largest Dashing experiment used 645 CPU hours on 100k genomes where NIQKI used only 30 CPU hours. We extrapolate that Dashing or Mash would need more than ten thousand CPU hours to handle 500k genomes while NIQKI used 166 CPU hours. On the downside, because of its large overhead NIQKI uses more memory than the other approaches on small collections and is slightly more memory expensive than Dashing on larger databases.

**Figure 7.**
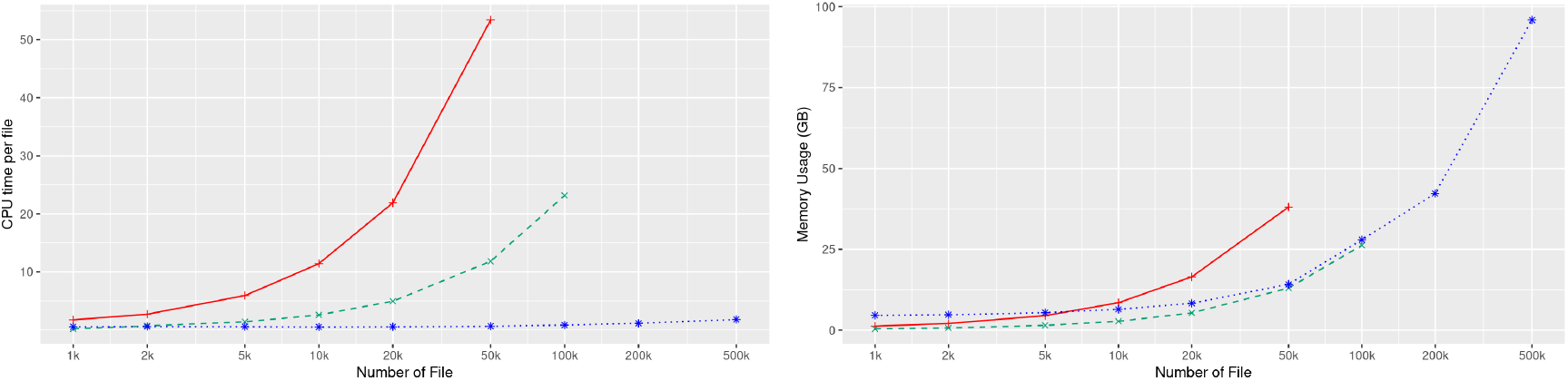
Benchmark on GenBank bacterial genomes. We report the CPU time divided by the number of entry and the total memory footprint for various collection sizes. Mash is plot in red, Dashing in green and NIQKI in blue.

### Indexing Genbank bacterial genomes

To access our approach scaling ability scalability, we choose to index and compute pairwise distance on all bacterial genomes from GenBank. This dataset represents more than one million genomes, counting more than five tera-nucleotides or more than one terabytes of gzipped fasta files. We choose to evaluate the cost of such an operation by varying the number of indexed minimizers that linearly impact our approach’s memory cost and the running time. We choose to keep the memory overhead constant 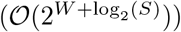 to ensure a fair memory comparison and raise the fingerprint size accordingly. We report the CPU time and Wall clock time along with the memory usage required for those experiments in Table 1. We observe that we can index and compute pairwise distance on such a database with medium-sized sketches in a few days with a reasonable memory footprint. If the indexing time is roughly constant and dominated by reading the input, we observe that the query time grows linearly with the sketch size (plus a “reading” time constant cost). We want to recall that most of the query computational time is due to the database’s redundancy, which generates many matches for some queries. For comparison, on a simulated dataset of one million random synthetics genomes indexed with 4096 fingerprints, the query time lasted 12 hours instead of 38 hours for the Genbank database.

**Table 1.** Benchmark on all Genbank bacterial genomes with various sketch sizes.

| #Fingerprint | CPU time (seconds) | Wall clock time all / query (hours) | Memory |
| --- | --- | --- | --- |
| 32 | 416,245 | 12 / 6 | 4,675 |
| 128 | 469,072 | 12 / 6 | 5,363 |
| 1,024 | 969,884 | 17 / 11 | 11,306 |
| 2,048 | 1,525,982 | 28 / 22 | 17,602 |
| 4,096 | 3,383,009 | 44 / 38 | 30,021 |

## Conclusions and future work

We showed that using inverted indexes on partitioned sketches leads to algorithmic improvement of fingerprint queries that can reduce running time by orders of magnitude with comparable precision. Theoretically, our proposed structure query is 𝒪 (#*hits*) compared to 𝒪(*S*.*N*) for state-of-the-art. This structure came with a memory cost as our index uses 𝒪(*S*(*NlogN* + 2^*W*^)) bits instead of 𝒪(*S*.*N*.*W*). We showed that even a straightforward implementation could efficiently index small fingerprints while providing results comparable to the state-of-the-art. Our approach could provide orders of magnitude faster queries on idealistic synthetic databases and real-world redundant databases with comparable memory footprints on large instances. We also demonstrated our index capacity to index and query all bacterial genomes of Genbank (more than one million bacterial genomes) in a matter of days on a small cluster. Our index can be used to index large collections to detect matches between novel query sequences and elements of the collection or to compute pairwise comparisons of all indexed sequences.

While our index can index any fingerprint (Minhash, Hyperloglog, Hyperminhash), we aimed to provide the best possible fingerprints of a given size to limit the number of false positives. To do so, we generalized the concept of Hyperminhash to account for different sizes of Hyperloglog and Minhash fingerprints dubbed (*h, m*)-HMH fingerprints. Interestingly Minhash, Hyperloglog, and Hyperminhash fingerprints can be seen as particular cases of (*h, m*)-HMH fingerprints. Given an expected sub-sampling, we can select the (*h, m*)-HMH fingerprint with the optimal parameter that provides the lowest false positive rate and confirms this choice in practice. While improving false-positive rates, those well-designed fingerprints come without computational or memory costs and could be used to improve existing sketching methods. Sketching methods could either use smaller fingerprints for a desired false positive rate, reducing memory footprint, or reducing their false positive rate without memory or time overhead.

On the practical aspect, our implementation still misses user-friendly features such as computing various metrics used in practice (Mash Index, containment index, or cardinality estimation) using advanced estimation methods. The different partition-based optimization described in the methods could also optimize query time in certain situations. Our running time could be improved by implementing classical optimization techniques such as batched queries, SIMD parallelism or vectorization, or practical optimization as computing only upper-half identifiers when comparing the index against itself. We mentioned that processing partition by column grants the possibility to stop queries that can not find high-scoring matches, but this feature is not yet implemented in practice. More generally more advanced output filtering technique could benefit our implementation in practice. More importantly, our current implementation uses plain integers as genomes identifiers leading to high memory costs. As our vectors can be seen as lists of increasing integers, those could be compressed with delta-encoding or other high throughput compression technique [8]. The index representation could be highly reduced with a limited impact on the runtime. While our proof of concept implementation is efficient, it tends to allocate a large amount of unused memory because of the heavy use of vectors. Other implementations could be made with different time/memory trade-offs. For example, comparing our approach with a Rank and the select-based index could be interesting.

On the theoretical aspects, our analysis of (*h, m*)-HMH fingerprint leads us to identify the need for an efficient fingerprint that presents smoother patterns to enable a larger range of possible hashes with or without estimation of the sub-sampling. On the fingerprint indexing problem, we showed that our partition query could be considered optimal as its running time is linear with the size of the output. However, the classical usage is to ask for matches with a number of hits above a certain threshold. It would be interesting to investigate which indexes or algorithmic solutions would provide an optimal answer to this problem.

## Acknowledgements

We want to thank Camille Marchet, Pierre Doignies, organizers and participants of the Bioinformatics: from Algorithms to Applications conference, for their support and discussions on this project. The ANR SEQdigger supported this work.

## Footnotes

3 https://www.ncbi.nlm.nih.gov/genbank/statistics/

4 https://www.ncbi.nlm.nih.gov/genome/doc/ftpfaq/

